# Predation risk reweights reward preference through coordinated amygdala-prelimbic dynamics

**DOI:** 10.64898/2026.09.02.748813

**Authors:** Eun Joo Kim, Ji Hoon Jeong, June-Seek Choi, Jeansok J. Kim

**Affiliations:** Department of Psychology, University of Washington, Seattle, WA 98195-1525, USA; School of Psychology, Korea University, Seoul, 02841, Republic of Korea; Neuroscience Program, University of Washington, Seattle, WA 98195-1525, USA

## Abstract

A fundamental challenge for animals and humans is resolving competing survival demands under naturalistic threat, yet circuit-level mechanisms remain poorly understood. We developed a paradigm recapitulating a predator-prey encounter: Long-Evans rats emerged from a nest to forage in an open arena, choosing between preferred and standard reward locations while facing a robotic predator. We simultaneously recorded single-unit activity from the basolateral amygdala (BLA) and the prelimbic cortex (PL). Rats shifted to the safer option under threat. BLA neurons responded predominantly to the predator, whereas PL neurons encoded reward value, threat context, and behavioral state. Population decoding revealed threat encoding in BLA and multiplexed representations in PL. Threat-responsive BLA neurons were preferentially recruited into BLA-PL synchrony before safe choices, and information flow became biased from BLA to PL following predator attack. These findings identify coordinated corticolimbic dynamics as a candidate mechanism through which threat reweights reward-guided action selection under ecological risk.

## Introduction

Foraging provides a canonical reward-threat conflict: as predation risk increases, animals routinely shift from high-yield, exposed resource patches to safer, lower-return alternatives [1]. Although behavioral ecology has formalized these adaptive decision rules [2], the neural computations that dynamically integrate reward value with environmental threat to guide flexible action selection remain poorly defined.

Corticolimbic circuitry, particularly the reciprocal interactions between the basolateral amygdala (BLA, comprising the lateral and basal nuclei) and the medial prefrontal cortex (mPFC), is implicated in valuation, threat processing, and behavioral flexibility [3–13]. The BLA encodes threat-predictive stimuli [14–17], whereas the mPFC integrates contextual and motivational information to bias action selection [7, 18–24]. These complementary functions position the BLA–mPFC circuit as a candidate for computing tradeoffs between reward and danger during ongoing behavior.

Much of our understanding of this circuitry’s function derives from Pavlovian fear conditioning paradigms [25, 26]. Within this framework, BLA neurons exhibit transient increases in firing to a conditioned stimulus (CS; e.g., a tone) following CS-unconditioned stimulus (US; e.g., a footshock) pairings [17, 27], while prelimbic (PL) neurons in the mPFC display sustained activity correlated with conditioned freezing behavior [26]. Dual-region recordings in animals trained with footshock- and reward-predictive CSs have revealed directional BLA→PL coordination during compound CS presentations that predicts whether animals will freeze or pursue a reward; optogenetic and chemogenetic manipulations of this projection bidirectionally modulated defensive and appetitive behavior [25]. These findings demonstrate that BLA→PL signaling can bias behavior when discrete cues signal competing motivational outcomes. However, cue-based paradigms provide limited insight into how this circuit supports ongoing value updating under naturalistic threat, as discrete CS-US pairings do not require the animal to continually re-evaluate reward value as risk fluctuates [28, 29].

In natural environments, threat is graded, continuous, and spatially structured, demanding ongoing cost-benefit computation rather than discrete conditioned responding [30]. Using a robotic predator paradigm in which rats foraged under ongoing predation risk [31, 32], Kim et al. [23] showed that BLA neurons exhibited phasic threat responses, whereas PL neurons displayed sustained activity unrelated to freezing.

Crucially, inter-regional synchrony increased during predator-relevant behaviors, indicating that corticolimbic circuits encode both internal threat states and real-time behavioral demands; however, that paradigm offered a single food source, so reward value itself was never placed in competition with threat. It therefore remains unknown how BLA–PL interactions dynamically reweight competing reward options as threat fluctuates—the central computation required for adaptive foraging under risk. This problem extends beyond predator avoidance. Across species and decision domains, adaptive behavior requires reweighting competing options as environmental risk changes—for example, when prey animals shift from high-yield to safer, lower-yield foraging patches under predation pressure [33–35] or humans reallocate capital from volatile assets to safe havens during market uncertainty [36–38]. Two key questions therefore remain unresolved: (i) how single neurons and neural populations within BLA and PL encode threat context (safe vs. risky), reward value (preferred vs. nonpreferred), and behavioral choice (stay vs. switch); and (ii) whether PL neurons multiplex these variables—representing threat, value, and choice within the same population—to support flexible action selection.

Here, we adapted the robotic-predator paradigm described above to directly pit competing food values against predatory threat, by introducing two food sources of unequal value. Simultaneous single-unit recordings in BLA and PL revealed coordinated, threat-dependent circuit dynamics that preceded shifts in reward preference.

Specifically, threat-encoding BLA neurons were selectively recruited into synchrony before safe choices, whereas directed BLA→PL information transfer emerged following active predator attacks. At the population level, decoding analyses revealed preferential threat encoding in the BLA, whereas PL maintained multiplexed representations of threat, reward value, and impending behavioral choice. Together, these findings suggest a circuit-level computational basis by which threat dynamically reweights reward preferences during adaptive decision-making under ecological risk.

## Results

### Conditional predatory threat shifts foraging preference

Rats underwent habituation (to acclimate to the nest), baseline (to establish individual foraging preference), and testing, which consisted of pre-robot (without robot) and robot sessions (Fig 1A, S1A Fig). Rats reliably chose the preferred (*P*) chocolate pellet over the non-preferred (*NP*) standard pellet during both the baseline and pre-robot sessions, indicating that individual pellet preferences remained stable before robot exposure. When predatory threat was introduced via Robogator surges triggered by approach to the *P* pellet, rats shifted their choices toward the *NP* pellet (Fig 1B). Relative to pre-robot sessions, rats exhibited a significantly lower preference index (PI; Fig 1C) and a greater number of failed *P* pellet trials (Fig 1D). These findings indicate that rats adaptively shifted their foraging preference away from the predator-associated high value reward toward the safer, lower-value alternative.

**Fig 1.**
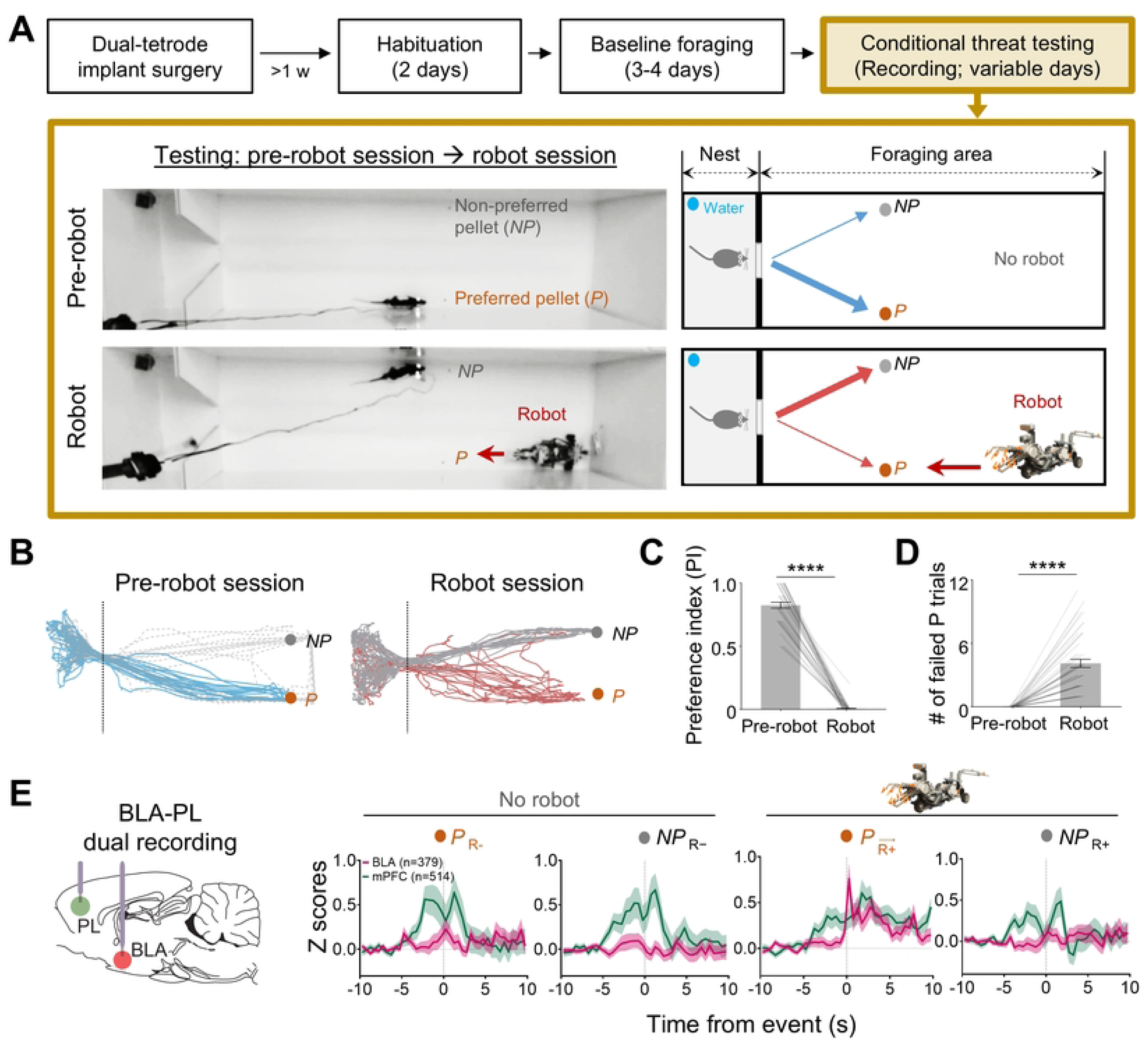
Conditional predatory threat shifts foraging preference and engages distinct BLA and PL population activity. **(A)** Experimental design. Rats with dual tetrodes targeting BLA and PL underwent nest habituation, foraging preference baseline, and conditional robot encounter testing. During the pre-robot session, animals chose between a preferred (*P*; chocolate) and a nonpreferred (*NP*; standard) pellet. During the robot session, robot activation was triggered by approach to the *P* pellet, whereas the robot remained stationary during approach to the *NP* pellet. **(B)** Representative foraging trajectories during pre-robot and robot sessions. Solid blue lines indicate initial trajectories when both pellet options were available; dashed gray lines indicate subsequent trajectories after *P* pellet had been obtained, leaving only the *NP* pellet available. Red lines indicate aborted approaches toward the *P* pellet following robot activation; solid gray lines indicate successful approaches to the *NP* pellet. **(C)** Preference index (PI) for the *P* pellet (n = 8 rats; 49 sessions). **(D)** Number of failed P pellet attempts (n = 8 rats; 49 sessions). **(E)** Population-averaged peri-event time histograms (PETHs) for BLA (n = 379) and PL (n = 514) neurons across four task events. Data are presented as mean ± SEM. \*\*\*\**P* < 0.0001. Detailed statistical results are provided in S1 Table.

### BLA neurons predominantly encode predatory threat whereas PL neurons represent multiple task variables

Histological verification confirmed tetrode placements within BLA and PL in all included animals (S1B and S1C Fig). In total, 379 BLA and 514 PL neurons were recorded across 8 animals. Representative single-unit recordings confirmed stable cluster isolation in both regions (S1C Fig).

Peri-event time histograms (Fig 1E) revealed that BLA and PL displayed distinct response profiles across four task events: preferred pellet without robot (*P* _R-_), nonpreferred pellet without robot (*NP* _R-_), preferred pellet with a surging robot (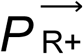), and nonpreferred pellet with a stationary robot (*NP* _R+_). Neural activity was aligned to pellet location entry for *P* _R-_, *NP* _R-_, and *NP* _R+_, and to robot activation for 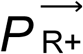.

BLA activity was dominated by a strong, time-locked response to the robot surge during *P* approach, with only minor modulation across the other three events (S1D Fig, left). In contrast, PL activity was broadly distributed across all four events, modulated by both reward value (pellet preference) and threat context (S1D Fig, right). Thus, BLA selectively signals dynamic threat, whereas PL multiplexes reward and contextual information.

Event-responsive neurons were classified into five functional types based on their excitation profiles (Fig 2A): *P* _R-_ cells (responsive during preferred-pellet approach in safe context), *NP* _R-_cells (responsive during nonpreferred-pellet approach in safe context), 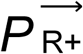 cells (responsive during the robot surge on preferred-pellet trials), *NP* _R+_ cells (responsive during nonpreferred-pellet approach in threat context), and multi-event (*M*) cells. *M* cells were further subdivided into *M2*, *M3*, and *M4* based on engagement across two, three, or four task events, respectively. Few cells showed event-related inhibition (S2A-S2G Fig); thus, analyses focused on event-excited cells, with inhibited and unmodulated cells grouped as *Others*. A greater proportion of PL than BLA neurons were task-responsive (Fig 2B). The distribution of single-versus multi-event responsiveness also differed markedly between regions: BLA neurons more often showed selective single-event responses (Fig 2C). BLA contained a significantly greater proportion of 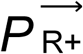 cells, whereas PL contained more *M* cells (Fig 2D). Within *M* cells, PL showed fewer *M2* and more *M3* cells than BLA, indicating a distinct shift toward broader event-conjunction coding in PL (Fig 2E).

**Fig 2.**
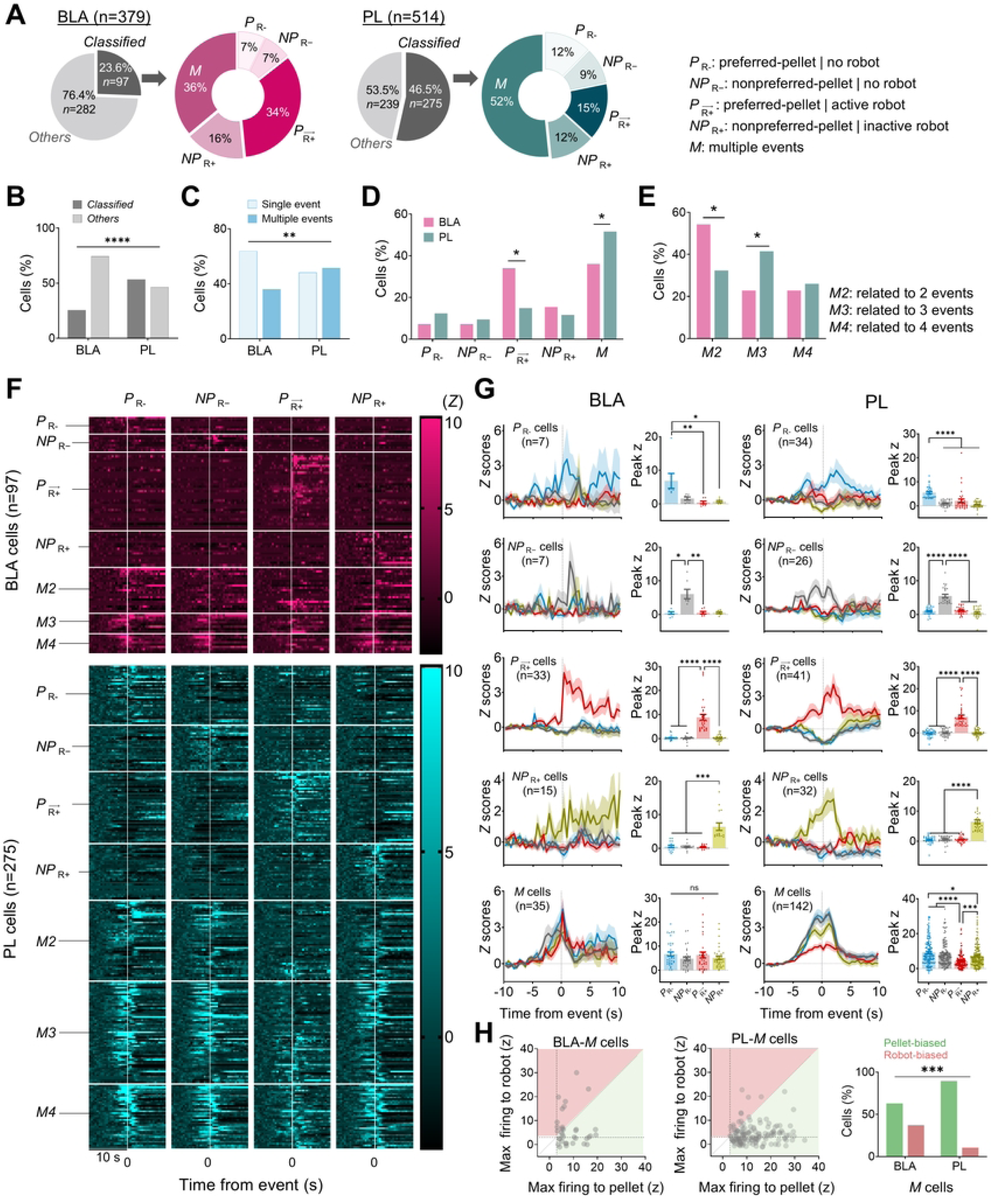
Functional organization of event-responsive neurons in BLA and PL. **(A)** Classification of event-responsive neurons. **(B)** Proportion of task-responsive neurons (BLA, n = 379 units; PL, n = 514 units). **(C)** Distribution of single-event- and multi-event-responsive neurons (BLA, 97 units; PL, 275 units). **(D)** Distribution of functional cell types (BLA, 97 units; PL, 275 units). **(E)** Distribution of *M2*, *M3*, and *M4* cells within the multi-event-responsive population (BLA, 35 units; PL, 142 units). **(F)** Color-coded PETHs sorted by response type. **(G)** Peak firing rates across task events for each functional cell type. **(H)** Robot-bias versus pellet-bias analysis of multi-event-responsive neurons (BLA, 35 units; PL, 142 units). Data are presented as mean ± SEM. \**P* < 0.05, \*\**P* < 0.01, \*\*\**P* < 0.001, \*\*\*\**P* < 0.0001. Detailed statistical results are provided in S1 Table.

Among single-event-responsive neurons (Fig 2F), peak firing rates to the corresponding events significantly exceeded those to non-corresponding events in both regions (Fig 2G; top four rows). However, the functional profile of multi-event neurons diverged sharply between structures. BLA *M* cells showed no significant differences in peak firing rates across events, which may reflect generalized salience coding. In contrast, PL *M* cells exhibited graded responses, with highest activity during pre-robot pellet procurement and lowest during the robot surge (Fig 2G; bottom row). BLA contained a greater proportion of robot-biased neurons, whereas PL contained a greater proportion of pellet-biased neurons (Fig 2H). Thus, BLA neurons are predominantly threat-selective and salience-driven, whereas PL neurons integrate reward value, threat context, and event structure—consistent with complementary roles in threat detection versus flexible decision-making.

### BLA–PL spike synchrony precedes choice under predatory threat

Given the distinct response profiles of the BLA and PL populations, we next examined which functional cell types were preferentially recruited into BLA-PL spike synchrony preceding behavioral choice. BLA-PL spike synchrony was quantified using cross-correlograms (CCs) during approach epochs preceding risky choices (*P* approach) and safe choices (*NP* approach) under predatory threat (Fig 3A-3B). Only sessions with at least three *P* attempts triggering robot surges were included, as *NP* success reduced repeated approaches to *P*.

**Fig 3.**
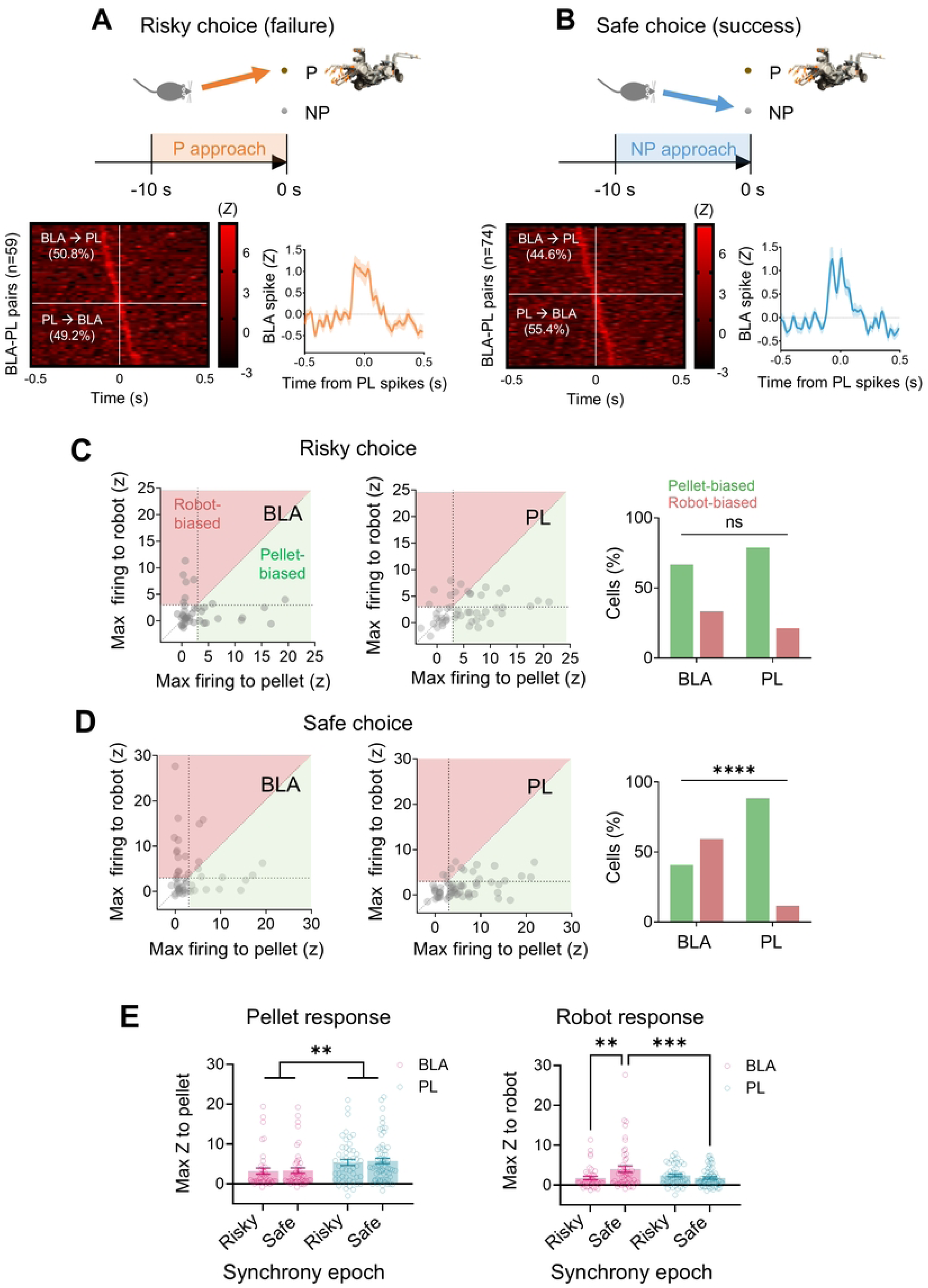
Functional composition of BLA-PL synchrony preceding risky and safe choices. **(A** and **B)** Significant BLA-PL spike synchrony during *P*-approach (risky-choice) and *NP*-approach (safe-choice) epochs. **(C** and **D)** Functional composition of synchronized pairs during risky (BLA, 13 units; PL, 33 units) and safe (BLA, 27 units; PL, 43 units) choices. **(E)** Pellet-related and robot-related responses of neurons participating in synchronized pairs (BLA, 41 units; PL, 50 units). Data are presented as mean ± SEM. \*\**P* < 0.01, \*\*\**P* < 0.001, \*\*\*\**P* < 0.0001. Detailed statistical results are provided in S1 Table.

A total of 1,971 BLA-PL neuron pairs were analyzed; 574 *P*-approach and 644 *NP*-approach pairs met the firing threshold for CC analysis > 0.1 Hz in both regions. Of these, 59 pairs exhibited significant spike synchrony during *P*-approach epochs (Fig 3A) and 74 pairs during *NP*-approach epochs (Fig 3B). The proportions of BLA-leading pairs (*P*-approach: 50.8%; *NP*-approach: 44.6%) and PL-leading pairs (*P*-approach: 49.2%; *NP*-approach: 55.4%) did not differ significantly between choice types.

During risky choices, the proportions of robot-biased versus pellet-biased neurons in synchronized pairs did not differ significantly between the BLA and PL (Fig 3C). In contrast, during safe choices, synchronous BLA–PL pairs were preferentially composed of robot-biased BLA cells and pellet-biased PL cells (Fig 3D). Consistent with this functional segregation, pellet-evoked firing was greater in synchronized PL neurons than in synchronized BLA neurons (Fig 3E). Conversely, robot-evoked responses were stronger in BLA neurons synchronized during safe choices, exceeding those observed in BLA neurons synchronized during risky choices and in synchronized PL neurons (Fig 3E). Moreover, BLA neurons participating in synchrony during safe choices, but not risky choices, exhibited stronger robot-surge responses than non-synchronized BLA neurons (S3A and S3B Fig). In contrast, synchronized and non-synchronized PL neurons did not differ in overall event-related activity, likely because elevated task-related activity was broadly distributed across the PL population (S3C Fig).

Together, these findings indicate that threat-selective BLA neurons are preferentially recruited into BLA–PL synchrony specifically when animals choose the safer option, whereas PL maintains value-related signals irrespective of choice.

### PL proactively encodes multiple task variables whereas BLA decoding is reactive and threat-locked

Linear support vector machine (SVM) classifiers were applied to four decoding contrasts (Fig 4A and 4B): (i) preference coding in the safe context, (ii) preference coding in the risky context, (iii) context coding for the preferred option, and (iv) context coding for the nonpreferred option.

**Fig 4.**
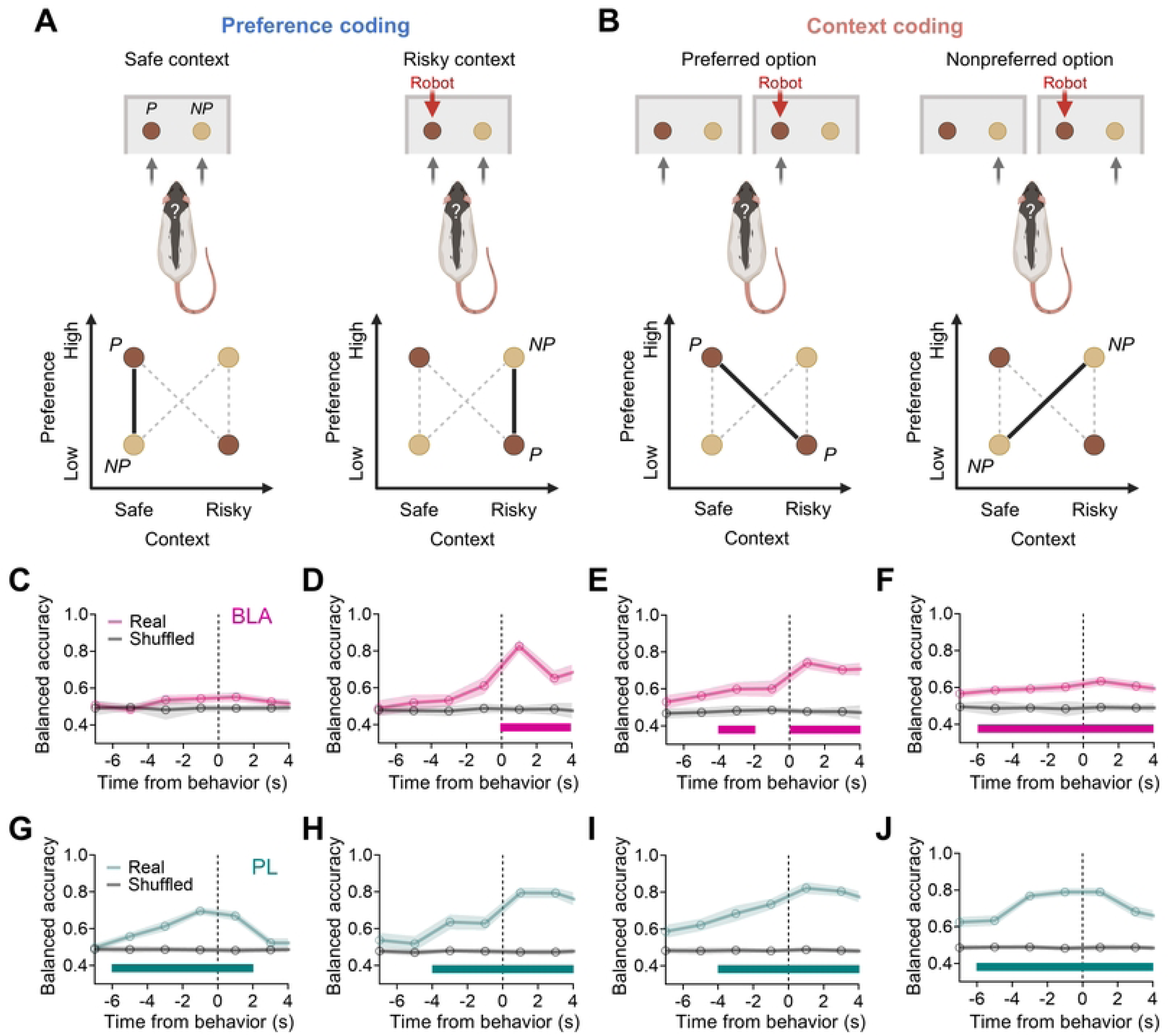
Population decoding of preference and context in BLA and PL. **(A)** Preference coding schematic. **(B)** Context coding schematic. (*C-F*) BLA decoding accuracy for preference (safe context: 39 sessions; risky context: 22 sessions) and context (preferred option: 22 sessions; nonpreferred option: 39 sessions) contrasts. (*G-J*) PL decoding accuracy for preference (safe context: 39 sessions; risky context: 22 sessions) and context (preferred option: 22 sessions; nonpreferred option: 39 sessions) contrasts. Shuffled controls are shown as mean ± SD. Horizontal bars indicate significant decoding relative to shuffled controls (*P* < 0.05). Detailed statistical results are provided in S1 Table.

BLA decoding capacity was limited for preference coding but more sustained for context coding. The BLA failed to decode pellet preference in the safe context (accuracy vs. shuffled; Fig 4C). Significant above-chance decoding was observed only in post-event windows (0-4 s) for preference in the risky context (Fig 4D). In contrast, context coding of the preferred option showed significant decoding across both pre- and post-event windows (Fig 4E). Context coding for the nonpreferred option showed sustained above-chance accuracy across an even broader temporal window (Fig 4F). In contrast, PL decoded all four contrasts with accuracy significantly above shuffled controls across all peri-event windows, including pre- and post-choice epochs (Fig 4G-4J). Decoding accuracy in PL was significantly greater than in BLA during pre-choice windows across all four contrasts (S4A-S4D Fig). Baseline decoding (−12 to −8 s before each event) did not differ from chance for either region across three of the four contrasts, except for context coding for the nonpreferred option, which showed above-chance accuracy at baseline in both BLA and PL (S5 Fig), suggesting that contextual information for this comparison is represented prior to task engagement.

These results indicate that PL maintains proactive, multiplexed representations of task variables available before choice execution, whereas BLA decoding is predominantly reactive, threat-locked, and temporally restricted.

### Directed information flow from BLA to PL increases selectively following predator attack

To characterize the directionality and temporal profile of BLA-PL communication, we applied an information-theoretic framework sensitive to both linear and nonlinear inter-regional dependencies. Each session was divided into five event epochs (Fig. 5*A*): control (inside the nest, pre-robot), *P* _R-_ and *NP* _R-_ (*P* and *NP* approach, pre-robot), 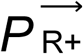 (*P* approach, active robot), post-surge (*P* retreat, post-active robot) and *NP* _R+_ (NP approach, inactive robot).

**Fig 5.**
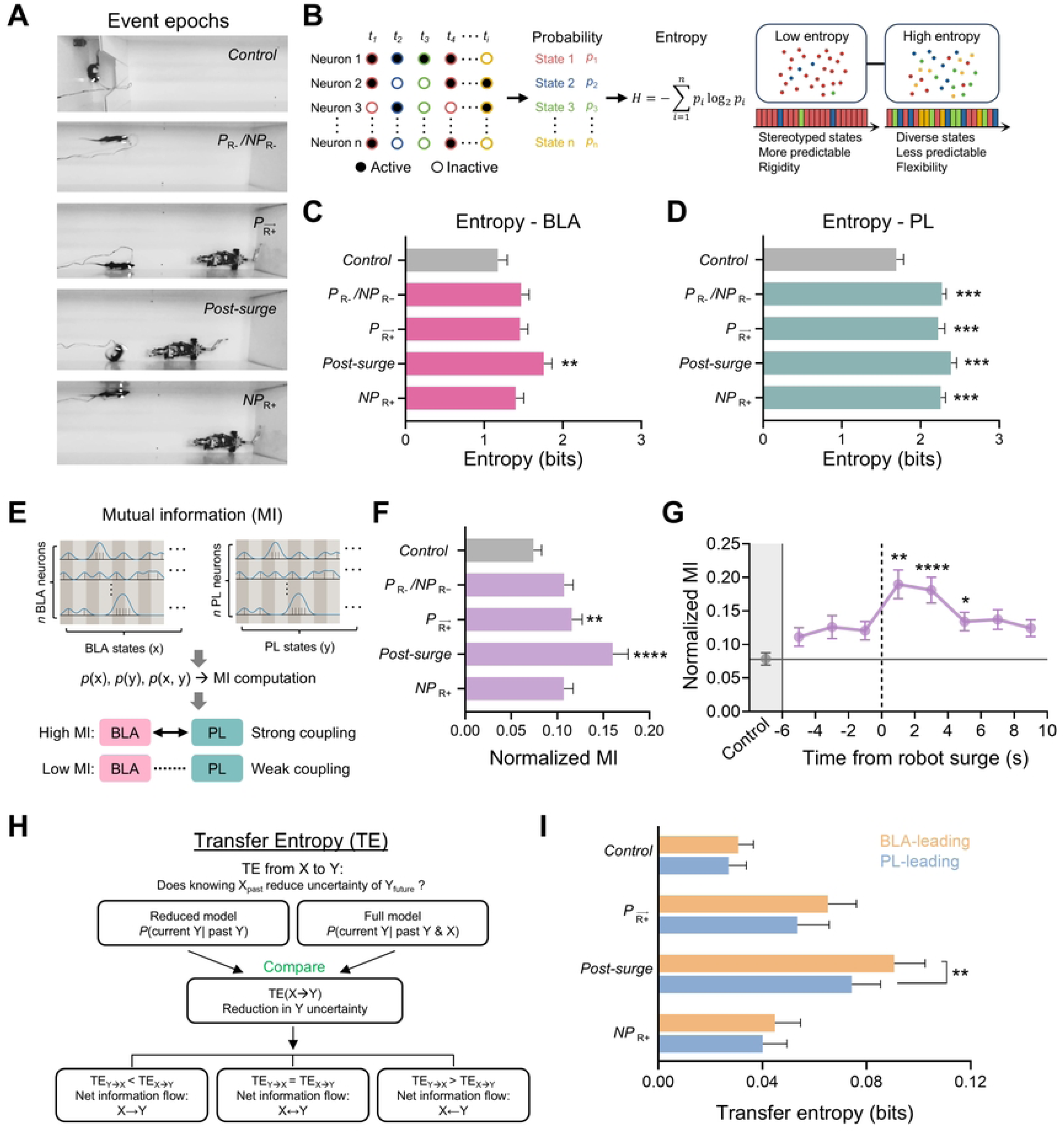
Entropy, mutual information, and transfer entropy analyses of BLA-PL communication. **(A)** Behavioral epochs used for information-theoretic analyses. **(B)** Entropy analysis schematic. **(C** and **D)** Shannon entropy across behavioral epochs for BLA and PL (39 sessions). **(E)** Mutual information (MI) analysis schematics. **(F)** Normalized BLA–PL mutual information across behavioral epochs (39 sessions). **(G)** Sliding-window mutual information aligned to robot activation (39 sessions). **(H)** Transfer entropy (TE) analysis schematic. **(I)** Bidirectional transfer entropy across behavioral epochs (21 sessions). Data are presented as mean ± SEM. \**P* < 0.05, \*\**P* < 0.01, \*\*\**P* < 0.001. Detailed statistical results are provided in S1 Table.

Shannon entropy of population state distributions was computed for each region and epoch to assess whether population complexity varied across behavioral contexts (Fig 5B). In BLA, entropy increased significantly only during the post-surge epoch relative to control (Fig 5C). In contrast, PL entropy was elevated across all task-related epochs relative to control, with no significant differences among those epochs (Fig 5D). PL entropy was significantly higher than BLA across all epochs (S6 Fig). Thus, BLA population complexity increased selectively following predator attack, whereas PL complexity remained elevated throughout task engagement.

We next quantified functional coupling between BLA and PL using mutual information (MI), which quantifies the reduction in uncertainty about one region’s activity given knowledge of the other (Fig 5E and 5F). MI differed significantly across epochs.

Post-hoc analyses revealed MI was significantly elevated during 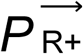 and post-surge epochs relative to control and *P* _R-_ and *NP* _R-_ epochs (Fig 5F); no other pairwise differences were significant. A sliding-window MI analysis further showed that MI increased sharply following the robot surge, peaked within 0-4 s, and remained elevated through +6 s (Fig 5G).

Although MI captures statistical dependence between neural populations without requiring model assumptions, it is symmetric and does not indicate directionality. To address this limitation, we computed transfer entropy (TE) to quantify directed information flow between regions (Fig 5H) [39–42]. TE was computed bidirectionally across four epochs. A repeated-measures two-way ANOVA with direction (BLA→PL vs. PL→BLA) and epoch as factors revealed significant main effects of both factors but no interaction. Post-hoc analyses showed directional asymmetry selectively during post-surge, where TE(BLA→PL) significantly exceeded TE(PL→BLA) (Fig 5I); directional asymmetry was absent in all other epochs.

Together, these analyses reveal a transient reorganization of BLA-PL communication following predator attack. BLA population complexity increased selectively during the post-surge epoch, interregional coupling rose sharply around the robot surge, and information flow became preferentially directed from BLA to PL. These findings indicate that predator activation is associated with enhanced and directionally biased corticolimbic communication.

### Robot-responsive BLA neurons and multi-event-responsive PL neurons are recruited into post-surge BLA-PL synchrony

To determine whether enhanced BLA-PL coupling during predatory threat was reflected in transient spike synchrony between simultaneously recorded pairs, we next examined whether transient spike synchrony between simultaneously recorded BLA-PL pairs varied across robot surge epochs and whether specific functional cell types were differentially recruited. CCs were computed in 4-s windows immediately preceding (pre-surge) and following (post-surge) robot activation (Fig 6A) [23]. Of 1,971 BLA-PL pairs analyzed, 418 pre-surge and 517 post-surge pairs met the firing threshold for CC analysis (> 0.1 Hz in both regions). Of these, 57 pairs exhibited significant spike synchrony during the pre-surge epoch (Fig 6B and 6D), whereas 79 pairs exhibited significant synchrony during the post-surge epoch (Fig 6C and 6F); only 9 pairs met criteria in both epochs.

**Fig 6.**
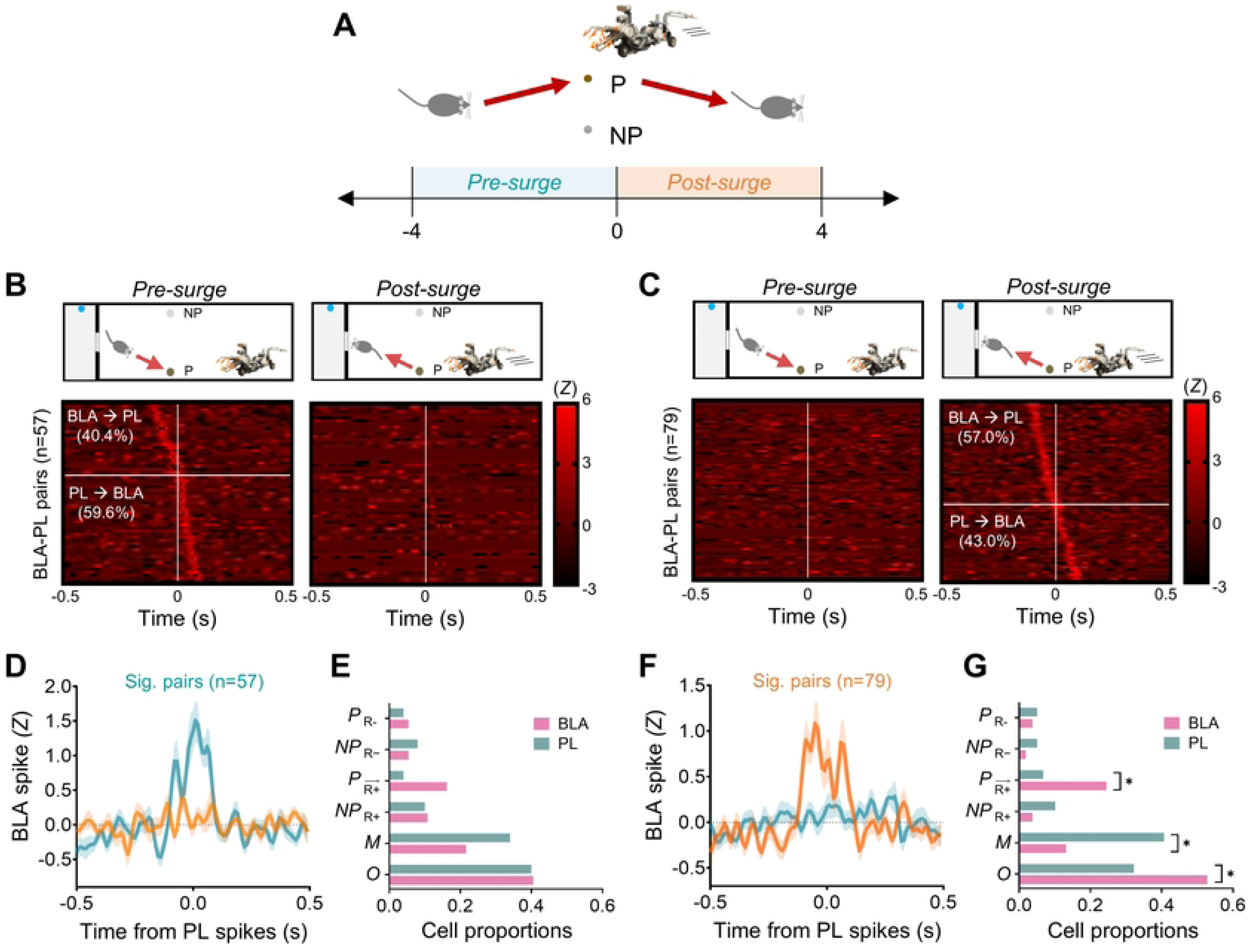
Functional composition of synchronized BLA-PL ensembles before and after robot activation. **(A)** Pre-surge and post-surge analysis windows. **(B** and **C)** Significant BLA-PL cross-correlograms during pre-surge and post-surge epochs. **(D** and **F)** Mean cross-correlograms of significant synchronized pairs. **(E** and **G)** Functional cell-type composition of synchronized pairs during pre-surge and post-surge epochs (BLA, n = 37 units; PL, n = 50 units). Data are presented as mean ± SEM. \**P* < 0.05. Detailed statistical results are provided in S1 Table.

The distribution of leading interactions differed across epochs. PL-leading interactions accounted for 59.6% of synchronized pairs during the pre-surge epoch, whereas BLA-leading interactions accounted for 57.0% during the post-surge epoch (Fig 6B and 6C). During the pre-surge epoch, the proportions of functional cell types represented in synchronized pairs did not differ between BLA and PL (Fig 6E). In contrast, during the post-surge epoch, robot-responsive BLA neurons and multi-event-responsive PL neurons were disproportionately represented among synchronized pairs (Fig 6G). The proportion of non-responsive (*Others*) cells also differed significantly between regions, with PL contributing a greater proportion than BLA.

Together, these findings show that the cellular composition of synchronized BLA-PL ensembles differed across robot surge epochs. Post-surge synchronized ensembles were enriched for robot-responsive BLA neurons and multi-event-responsive PL neurons, while the prevalence of leading interactions shifted from PL-leading before the surge to BLA-leading after the surge. Consistent with the post-surge increase in BLA→PL transfer entropy, these findings suggest that threat-responsive BLA neurons and multi-event-responsive PL neurons contribute disproportionately to corticolimbic coordination following predator activation.

## Discussion

The present findings suggest that naturalistic predatory threat is associated with flexible reward preference reweighting and complementary coding across a corticolimbic circuit. The results are consistent with a model in which threat-related signals in BLA interact with multiplexed representations in PL to support adaptive action selection under ecological risk. This work extends corticolimbic circuit function beyond discrete conditioning paradigms to continuous value-based decision-making in a naturalistic setting.

Optimal foraging theory predicts that animals abandon high-yield but risky patches when predation risk exceeds a threshold, shifting to safer alternatives to balance energetic gain against predation risk [1],[2]. Our behavioral findings align with this framework: rats shifted preference from a high-value to a lower-value option when threat was contingently imposed at the preferred location while continuing to forage actively. Failed approaches to the preferred option increased during robot sessions, consistent with graded, proximity-dependent risk assessment described in predatory imminence frameworks [43–45] rather than generalized avoidance. Unlike paradigms employing discrete conditioned cue presentations within structured trials, the present task required continuous evaluation of competing reward and threat values. The persistence of foraging despite threat exposure indicates that predatory risk reweights reward preference rather than suppresses reward seeking, highlighting a neural process that may support adaptive choice under uncertainty.

The dissociation between threat-biased BLA coding and multiplexed PL coding is consistent with a mixed selectivity framework [22, 46–48], in which flexible behavior emerges from neural representations that encode combinations of task-relevant variables. A recent perspective distinguishes pure selectivity, linear mixed selectivity, and nonlinear mixed selectivity, with nonlinear mixed selectivity increasing representational dimensionality and enabling flexible readout by downstream circuits [48]. The present findings provide empirical support for this conceptual synthesis, with BLA and PL occupying dissociable positions along the proposed coding continuum. BLA 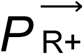 cells, with their selective, time-locked responses to the predator, are consistent with a specialized threat-detection function. In contrast, PL *M* cells exhibited context-dependent modulation across reward value, threat, and behavioral state, consistent with nonlinear mixed selectivity. This distinction may explain why PL—but not BLA—carried anticipatory representations of both reward preference and threat context prior to behavioral transitions.

Notably, PL contained a higher proportion of task-responsive neurons than BLA (53.5% vs. 23.6%), with responses of broader task coverage, consistent with the proliferative coding strategy attributed to the prefrontal cortex [48]. This contrasts with Pavlovian fear conditioning studies, in which PL activity has been primarily associated with conditioned freezing [26, 49]; under predatory threat, PL activity is not linked to freezing but instead reflects heterogeneous encoding of reward value, threat, and behavioral state. Although PL lesions have previously been shown to abolish context-appropriate fear expression in cue-conditioned paradigms [20], the present findings extend this context sensitivity to the continuous integration of multiple task-relevant variables during naturalistic foraging. This capacity for context-dependent reconfiguration is consistent with recent evidence that mPFC ensembles dynamically switch between goal-directed and threat-avoidance coding modes during foraging under robotic predator threat [50], suggesting adaptive coding according to reward-threat contingencies. A similar principle of functional specialization has been observed at the single-cell level in BLA, where fear conditioning recruits distinct fear-expression and extinction populations whose balance predicts behavioral state transitions [14]; the present robot-responsive cell population may represent an analogous threat-selective BLA subpopulation specialized for predator encounters.

The data further suggest a division of labor between threat detection and value integration. BLA activity was predominantly threat-locked, whereas PL showed broader engagement across task variables and behavioral epochs. A complementary dissociation has been identified in humans using virtual predator paradigms, where distal threat engages ventromedial prefrontal cortex for cost-benefit evaluation while proximal threat shifts activity to the periaqueductal gray for reflexive defense [51]; the reactive BLA and proactive PL profiles observed here may reflect an analogous segregation of threat-reactive and threat-evaluative processes within the rodent corticolimbic circuit.

Interregional synchrony and information-transfer analyses further indicate that communication between the two regions becomes selectively enhanced around predator encounters, suggesting that threat-related amygdala signals update ongoing prefrontal representations as environmental risk changes. Prior work in fear conditioning has shown that 4-Hz oscillations and theta-gamma coupling in the BLA-PL circuit increase during fear expression to conditioned stimuli [52, 53]; the present data extend this observation to naturalistic foraging, suggesting that BLA→PL coordination during threat-driven behavioral transitions is not limited to conditioned stimulus presentations but may reflect a general mechanism for adaptive reconfiguration of prefrontal representations. Rather than functioning as a simple feedforward threat pathway, the BLA-PL circuit may support dynamic interactions between specialized and multiplexed representations, enabling behavioral preferences to be adjusted as predation risk changes.

These observations motivate a circuit model of adaptive preference reweighting. In this framework, predator encounters are associated with transient BLA activity that temporally precedes increased BLA–PL coordination. Such dynamics may permit behavioral priorities to shift as environmental risk evolves, providing a mechanism through which animals balance resource acquisition against survival demands. Although the present findings are correlational, they generate testable predictions that threat-responsive BLA populations and multi-event PL populations play causal roles in behavioral preference reweighting.

Beyond foraging, the principles identified here may generalize to broader threat-reward conflicts. Under normative conditions, threat signals must bias behavior sufficiently to avoid danger without excessively suppressing reward pursuit. Supporting this view, individuals with focal bilateral amygdala lesions caused by Urbach-Wiethe disease show diminished monetary loss aversion, failing to appropriately downweight risky options when potential losses are present [54]. This parallels our finding that BLA threat encoding was associated with preference reweighting away from a threatened higher-value option, suggesting a conserved role for the amygdala in loss-sensitive decision-making across species and contexts. Disruption of this regulatory balance— through exaggerated amygdala signaling, impaired prefrontal integration, or disrupted interregional communication—could contribute to maladaptive avoidance and threat generalization, consistent with circuit models of post-traumatic stress disorder and related anxiety disorders [55–58]. The present paradigm may provide an ethologically grounded framework for investigating how corticolimbic circuits regulate behavioral responses across adaptive and maladaptive states.

Several limitations warrant consideration. Recordings were restricted to male animals, and whether similar dynamics occur in females remains unknown given established sex differences in amygdala-dependent threat processing [59, 60]. In addition, our analyses are correlational and therefore cannot establish causal contributions of BLA-PL interactions to behavioral switching. Transfer entropy was estimated from a discretized representation of neural population activity. Although the coarse state representation was necessary for reliable probability estimation, it likely underestimated the magnitude of directed information transfer between BLA and PL. Finally, downstream targets of PL, including the basal ganglia, nucleus reuniens, and hypothalamic structures [61–66], were not examined and are likely to contribute to translating corticolimbic computations into behavior.

The present findings identify complementary coding and coordinated communication within a corticolimbic circuit during adaptive foraging under threat. By associating optimal foraging behavior with neural mechanisms of threat-reward integration, this work provides a framework for understanding how survival demands reshape value-based decision-making in dynamic environments.

## Materials and Methods

### Animals

Adult male Long–Evans rats (Charles River Laboratories) were used. Animals were singly housed on a 12 h light/dark cycle with ad libitum access to water. During behavioral training and testing, rats were food restricted to ∼85% of their free-feeding body weight. All experiments were conducted during the dark phase. Procedures were approved by the University of Washington Institutional Animal Care and Use Committee (IACUC protocol #4040-01) and conformed to NIH guidelines.

### Surgery

Rats were anesthetized with isoflurane (1–2%) and secured in a stereotaxic frame (Kopf Instruments). A custom microdrive array containing independently movable tetrodes was implanted unilaterally targeting the BLA (AP −2.8 mm, ML +5.0 mm; 6 tetrodes) and PL (AP +3.0 mm, ML +0.7 mm; 6 tetrodes), relative to bregma. Tetrodes were constructed from 14 μm formvar-insulated nichrome wire (Kanthal) and gold-plated to a final impedance of 100–300 kΩ measured at 1 kHz. The microdrive assembly was secured to the skull with bone screws and dental acrylic. Animals recovered for at least 7 days before behavioral training and recording.

### Behavioral Procedure

Rats were habituated to the apparatus by placement in the nest compartment for 30 min per day on two consecutive days. During habituation, standard and chocolate pellets (10 each; Bio-Serv, 500 mg) were available in the nest compartment along with water to familiarize animals with both pellet types and the testing environment.

Baseline foraging consisted of 10 trials per day for 3–4 days. In each trial, rats exited the nest and chose between two fixed foraging locations positioned equidistant from the nest. One location contained a chocolate pellet (preferred, *P*) and the other contained a standard pellet (non-preferred, *NP*); the spatial assignment of pellet type was counterbalanced across animals. After retrieval of the first pellet, rats returned to obtain the remaining pellet, completing the trial. Preference was defined as a significantly greater proportion of *P* choices across baseline sessions.

Robogator encounter testing consisted of two sequential phases conducted on the same day. During the pre-robot session (median: 10 trials; range: 9-13 trials), rats freely chose between *P* and *NP* pellets in the absence of robotic movement. Each trial ended after both pellets had been retrieved and returned to the nest. During the robot session (median: 10 trials; range: 6-15 trials), forward movement (“surge”) of the Robogator was triggered when the rat crossed a predefined proximity threshold near the preferred pellet, whereas the robot remained stationary during approaches to the non-preferred pellet.

Behavioral measures included pellet preference (Preference index for *P* vs. *NP*) and the number of failed *P* trials. Threat-induced preference switching was defined as a reduction in *P* choices during the robot session relative to the pre-robot session.

### Electrophysiological Recording

Neural activity was recorded using a Neuralynx Cheetah digital acquisition system. Signals were amplified (10,000×), bandpass filtered (spikes: 600–6,000 Hz; local field potentials [LFPs]: 0.1–1,000 Hz), and digitized at 32 kHz. Spike waveforms were threshold detected and stored for offline analysis. Single units were isolated using SpikeSort3D (Neuralynx) followed by manual cluster refinement based on waveform amplitude, energy, and principal component features across tetrode channels. Units were included if clusters were well separated, stable across the recording session, and exhibited clear refractory periods in autocorrelograms. Only sessions with stable simultaneous recordings from BLA and PL were included in inter-regional analyses. Peri-event time histograms (PETHs) and raster plots were generated using NeuroExplorer (Nex Technologies).

### Event-Related Firing Analysis

Spike trains were aligned to behavioral events including nest exit, pellet approach, Robogator surge onset, and pellet retrieval. PETHs were constructed using 0.5 s bins[23]. Event-related modulation was assessed by comparing firing rates during defined event windows to baseline epochs (−10 to −5 s relative to each event) using z-normalization. Neurons showing significant modulation (Z > 3) within a 1.5 s window around a given event were classified as responsive to that event. Neurons responding to at least two events were classified as multiple event-related cells.

### Spike Synchrony and Cross-Correlation Analysis

Functional coupling between simultaneously recorded BLA and PL neurons was quantified using cross-correlation (CC) analysis with spike trains binned at 10-ms resolution. Shift predictors generated from 100 random trial shuffles were subtracted from the raw CCs to control for correlations arising from shared task structure. Shuffle-corrected correlograms were converted to Z-scores based on the mean and standard deviation of the shuffled distribution. Neuron pairs were considered significantly synchronized if peak Z-scores exceeded 3 within a ±100 ms window. Analyses were restricted to units with mean firing rates ≥ 0.1 Hz during the relevant behavioral epochs to avoid false peaks in the CCs. Changes in spike synchrony were assessed by comparing the following robot session epoch pairs: (i) *P* approach vs. *NP* approach and (ii) pre-surge vs. post-surge.

### Population Decoding Analysis

To assess whether population activity in BLA and PL encodes pellet preference and threat context, we trained linear support vector machine (SVM) classifiers on population neural data. Spike timestamps for each unit were converted into 1 kHz binary vectors, convolved with a Gaussian kernel (σ = 100 ms, width = 1000 ms), and z-scored using the mean and standard deviation calculated across the entire session. Sessions with fewer than three neurons in either region were excluded. Neural data from 2-s windows centered on the target event were extracted. The processed neural data were averaged into 100-ms bins. All data from the same region were concatenated into a single population vector. Linear SVM classifiers were implemented using the Python scikit-learn package. Input features were clipped to an absolute value of 5 to reduce the influence of outliers. To account for imbalanced trial counts across conditions, classification performance was evaluated using balanced accuracy with leave-one-out cross-validation. To establish a baseline, the same procedure was repeated with shuffled labels, and balanced accuracy was compared between true and shuffled conditions using repeated-measures ANOVA. Detailed parameters for the SVM classification analysis can be found in the publicly accessible code repository.

### Mutual Information Analysis

To assess inter-regional communication across behavioral contexts, we computed mutual information (MI) between BLA and PL population activity during five event epochs. All epoch windows for the events were set to 5 s in duration. The *control* epoch was centered on the midpoint between the last pre-robot approach and the first robot-session approach. *P* _R-_ and *NP* _R-_ epochs centered on pellet retrieval (−2.5 to +2.5 s) during the pre-robot session. 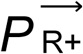 and *post-surge* epochs were defined as −5 to 0 s and 0 to +5 s relative to robot surge onset, respectively. The *NP* _R-_ epoch spanned −2.5 to +2.5 s relative to *NP* pellet retrieval during the robot session. To equalize sample sizes across epochs, the epoch type with the fewest events within each session was identified and all other epoch types were subsampled to match this count. The *control* epoch length was scaled accordingly to yield an equivalent number of bins. Neural data were processed as described above: spike trains were convolved with a Gaussian kernel (σ = 100 ms, width = 1000 ms), z-scored, and averaged into 50 ms bins. Population vectors were constructed by concatenating activity across neurons within each region for each time bin. Sessions with fewer than 3 units in either region were excluded. Population states were defined using k-means clustering (K = 10) applied to concatenated BLA and PL activity across all epochs, ensuring a shared state space. MI was calculated from the joint and marginal probability distributions of BLA and PL state labels and normalized by the geometric mean of individual region entropies. Shannon entropy was computed independently for each region and epoch. For the sliding-window analysis, data from −6 to +10 s relative to robot surge onset were segmented into non-overlapping 2 s windows, and MI was computed per window using the same approach.

### Transfer Entropy Analysis

To assess directionality of information flow between BLA and PL, we computed transfer entropy (TE) [39] using a discrete-state approach adapted from the MI pipeline. TE quantifies how much a source region’s past activity reduces uncertainty about a target region’s future state, above and beyond the target’s own history. A discrete-state estimator was used for methodological consistency with the MI analysis. Four behavioral epochs were analyzed: *control*, 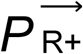, *post-surge*, and *NP* _R+_. Epoch duration was set to 6 s based on the MI temporal dynamics analysis. To equalize statistical power across epochs, the minimum event count within each session was identified, and all other epochs were subsampled to match this value. Subsampling was repeated 30 times, and TE values were averaged across iterations. Neural data preparation followed the MI pipeline with two modifications: the Gaussian kernel was narrowed (σ = 50 ms, width = 500 ms) to improve temporal resolution, and the number of k-means clusters was reduced to K = 5 to ensure sufficient sampling of state-transition. Clustering was performed on concatenated data across all epochs prior to subsampling to ensure consistent state assignments. Sessions with fewer than 3 neurons per region or fewer than 3 matched events per epoch were excluded. TE was computed in both directions (BLA→PL and PL→BLA), using a Markov order of k = 2 (100 ms history) and a prediction step of u = 1 (50 ms). TE was estimated using the four-entropy decomposition: *TE*_*J→I*_ = *H*(*i_t_*, *i_t-1_*) − *H*(*i_t-1_*) − *H*(*i_t_*, *i_t-1_*, *j_t-1_*) + *H*(*i_t-1_*, *j_t-1_*), where ‘*i*’ denotes the target region and ‘*j*’ the source region.

### Statistical Analyses

All statistical analyses were conducted using MATLAB (MathWorks), SPSS (v19), GraphPad Prism (v9.0), and NeuroExplorer (v5.030). All tests were two-tailed with significance set at α = 0.05. Data were assessed for normality prior to parametric testing; when assumptions were violated, nonparametric equivalents were used (Friedman test, Mann-Whitney test). Group comparisons used one-way or mixed-design ANOVA for within- and between-subject factors, with post hoc comparisons using Tukey’s HSD, Bonferroni, Dunn’s test, or Fisher’s LSD where appropriate. The Greenhouse–Geisser correction was applied when sphericity assumptions were violated. Paired or unpaired t-tests were used for planned comparisons, and Pearson’s correlation for linear associations, and chi-square tests followed by post hoc Z-tests for proportional comparisons. Sample sizes, exact p-values, and statistical details are reported in Table S1.

### Histology

At the conclusion of experiments, rats were deeply anesthetized and electrolytic lesions were made at tetrode tips by passing 10 μA current for 10 s to mark recording sites. Animals were transcardially perfused with 0.9% saline followed by 10% buffered formalin. Brains were extracted, post-fixed overnight at 4°C, and immersed in 30% sucrose until saturated. Coronal sections (40–50 μm) were cut and stained with cresyl violet to visualize cytoarchitecture and Prussian blue to identify iron deposits at electrolytic lesion sites. Tetrode placements were verified under light microscopy with reference to the Paxinos and Watson rat brain atlas [67]. Only animals with confirmed tetrode placements within BLA and PL were included in analyses.

## Acknowledgments

This study was supported by National Institutes of Health grant MH099073 (J.J.K.) and by the National Research Foundation of Korea (NRF) through grants funded by the Ministry of Science and ICT (MSIT, RS-2025-00517214) (J.-S.C.). We thank Dr. Jeiwon Cho for consultation during the initial stages of this project, and Jane Romani and Harry Boo for technical assistance with experiments, data collection, and analysis. Illustrations were created with BioRender.com.

## Author Contributions

Conceptualization, E.J.K., J.-S.C., and J.J.K.; data curation, E.J.K. and J.J.K.; formal analysis, E.J.K., J.H.J., J.-S.C., and J.J.K.; funding acquisition, J.-S.C. and J.J.K.; investigation, E.J.K. and J.J.K.; methodology, E.J.K., J.-S.C., and J.J.K.; project administration, J.J.K.; resources, J.-S.C. and J.J.K.; software, E.J.K. and J.H.J.; supervision, J.-S.C. and J.J.K.; validation, E.J.K., J.H.J., J.-S.C., and J.J.K.; visualization, E.J.K. and J.H.J.; writing – original draft, E.J.K. and J.J.K.; writing – review & editing, E.J.K., J.H.J., J.-S.C., and J.J.K.

## Captions for supporting information

**S1 Fig. Foraging apparatus and BLA–PL recording sites.**

**(A)** Schematic of the foraging arena with dimensions. The red arrow indicates the direction of the robotic predator surge. **(B)** Histological reconstructions of recording sites in the BLA (left) and PL (right). **(C)** Dual tetrode configuration for simultaneous BLA–PL recordings (top left), representative photomicrographs of electrode implants (bottom left), and representative single-unit waveforms and cluster plots from the BLA and PL (right). **(D)** Comparisons of baseline (−10 to −5 s) and event epoch (−2.5 to +2.5 s) firing rates for BLA (n = 379; left) and PL (n = 514; right) neurons across four task events. Data are presented as mean ± SEM. \**P* < 0.05, \*\*\**P* < 0.001. Detailed statistical results are provided in S1 Table.

**S2 Fig. Event-inhibited neurons in BLA and PL.**

**(A)** Proportions of non-responsive, event-excited, and event-inhibited neurons in the BLA (top) and PL (bottom), along with the functional cell-type breakdown of event-inhibited neurons (pie charts, right). *P* _R-_ (inhibited during preferred-pellet approach in safe context); *NP* _R-_(inhibited during nonpreferred-pellet approach in safe context); *P* _R+_ (inhibited during the robot surge on preferred-pellet trials); *NP* _R+_ (inhibited during nonpreferred-pellet approach in threat context); and *M* (inhibited during multiple task events: two, three, or four task events). **(B)** Color-coded PETHs of event-inhibited neurons in BLA (top) and PL (bottom), sorted by inhibition type. **(C)** PETHs and peak z-scores across task events for each event-inhibited cell type in the BLA (n = 18; left) and PL (n = 61; right), sorted by inhibition type. **(D)** Proportions of each event-inhibited cell type in the BLA (n = 18) and PL (n = 64). **(E)** Distribution of M2, M3, and M4 subtypes among multi-event-inhibited neurons (BLA, n = 10; PL, n = 34). **(F)** Minimum and maximum z-scored firing responses to the robot and pellet events for multi-event-inhibited neurons in the BLA (left) and PL (right). Green and pink shading indicate pellet- and robot-biased regions, respectively. **(G)** Proportions of pellet-biased and robot-biased neurons among multi-event-inhibited M cells in the BLA (n = 18) and PL (n = 61). Data are presented as mean ± SEM. \**P* < 0.05, \*\**P* < 0.01, \*\*\**P* < 0.001, \*\*\*\**P* < 0.0001. Detailed statistical results are provided in S1 Table.

**S3 Fig. Synchronized versus non-synchronized BLA and PL neurons during risky and safe choices.**

**(A)** PETHs and peak z-scores across task events for BLA neurons that participated in (correlated) or did not participate in (non-correlated) in BLA–PL synchrony during risky (top; correlated: n = 41, noncorrelated: n = 338) and safe (bottom; correlated: n = 50, noncorrelated: n = 329) choices. **(B)** PETHs and peak z-scores across task events for PL neurons participating and not participating in BLA–PL synchrony during risky (top; correlated: n = 49, noncorrelated: n = 465) and safe (bottom; correlated: n = 62, noncorrelated: n = 452) choices. Data are presented as mean ± SEM. \**P* < 0.05, \*\**P* < 0.01, \*\*\**P* < 0.001, \*\*\*\**P* < 0.0001. Black asterisks indicate comparisons between correlated and non-correlated neurons; gray asterisks indicate comparisons between events within non-correlated neurons; pink asterisks indicate comparisons between events within correlated neurons. Detailed statistical results are provided in S1 Table.

**S4 Fig. Differences in decoding accuracy between BLA and PL populations. (A–D)** Comparison of BLA and PL decoding accuracy for preference coding in the safe context (A; 39 sessions), preference coding in the risky context (B; 22 sessions), context coding for the preferred option (C; 22 sessions), and context coding for the non-preferred option (D; 39 sessions). Data are presented as mean ± SEM. Horizontal bars indicate time periods during which PL decoding accuracy was significantly greater than that of the BLA (*P* < 0.05). Detailed statistical results are provided in S1 Table.

**S5 Fig. Baseline decoding accuracy.** Balanced decoding accuracy during the baseline epoch (−12 to −8 s relative to each behavioral event) for BLA and PL populations across four decoding contrasts (39, 22, 22, and 39 sessions, left to right). Baseline decoding accuracy did not differ from shuffled controls for most comparisons, except for context coding of the nonpreferred option, which exhibited above-chance decoding accuracy in both regions. Data are presented as mean ± SEM. \*\*\**P* < 0.001, \*\*\*\**P* < 0.0001. Detailed statistical results are provided in S1 Table.

**S6 Fig. Differences in entropy between BLA and PL populations, related to Figure 5**. Comparison of BLA and PL entropy during each epoch: *Control, P* _R-_ */ NP* _R-_*, P* _R+_*, Post-surge,* and *NP* _R+_ (39 sessions). Data are presented as mean ± SEM. \*\*\**P* < 0.001, \*\*\*\**P* < 0.0001. Detailed statistical results are provided in S1 Table.

**S1 Table.** Detailed statistical results for main and supporting information figures.

**S1 Data.** Data underlying figures and analyses.

